# GSR-DB: a manually curated and optimised taxonomical database for 16S rRNA amplicon analysis

**DOI:** 10.1101/2023.04.19.537515

**Authors:** Leidy-Alejandra G. Molano, Sara Vega-Abellaneda, Chaysavanh Manichanh

## Abstract

Amplicon-based 16S ribosomal RNA sequencing remains the most widely used method to profile microbial communities, as a low-cost and low-complexity approach. Reference databases are a mainstay for taxonomic assignments, which typically rely on popular databases such as SILVA, Greengenes, GTDB, or RDP. However, the inconsistency of the nomenclature across databases, and the presence of shortcomings in the annotation of these databases are limiting the resolution of the analysis. To overcome these limitations, we created the GSR database (Greengenes, SILVA, and RDP database), an integrated and manually curated database for bacterial and archaeal 16S amplicon taxonomy analysis. Unlike previous integration approaches, this database creation pipeline includes a taxonomy unification step to ensure consistency in taxonomical annotations. The database was validated with three mock communities and two real datasets and compared with existing 16S databases such as Greengenes, GTDB, ITGDB, SILVA, RDP, and MetaSquare. Results showed that the GSR database enhances taxonomical annotations of 16S sequences, outperforming current 16S databases at the species level. The GSR database is available for full-length 16S sequences and the most commonly used hypervariable regions: V4, V1-V3, V3-V4, and V3-V5.

**IMPORTANCE:** Taxonomic assignments of microorganisms have long been hindered by inconsistent nomenclature and annotation issues in existing databases like SILVA, Greengenes, GTDB, or RDP. To overcome these issues, we created GSR-DB, accurate and comprehensive taxonomic annotations of 16S amplicon data. Unlike previous approaches, our innovative pipeline includes a unique taxonomy unification step, ensuring consistent and reliable annotations. Validated with mock communities and real datasets, GSR-DB outperforms existing databases in providing species-level resolution, making it a game-changer for microbiome studies. Moreover, GSR-DB is designed to be accessible to researchers with limited computational resources, making it a powerful tool for scientists across the board. Available for full-length 16S sequences and commonly used hypervariable regions, including V4, V1-V3, V3-V4, and V3-V5, GSR-DB is a go-to database for robust and accurate microbial taxonomy analysis.

## INTRODUCTION

16S rRNA gene amplicon-based (16S) sequencing is a widespread method that has profoundly impacted the human microbiome characterisation, revealing important insights into the complex interactions between microorganisms and their hosts. This approach has allowed us to associate altered microbial profiles with diseases, including gut-associated conditions, inflammatory bowel disease, metabolic diseases, and colorectal cancer (1,2).

16S analysis involves various upstream steps, including quality control, read trimming, and taxonomic classification. Previous studies have reported the impact of bioinformatic pipelines in the microbial profiling of biological samples, highlighting the importance of reference databases for taxonomic prediction (3). Currently, the most widely used databases are Greengenes (4), Genome Taxonomy database (GTDB) (5), SILVA (6), and Ribosomal Database Project (RDP) (7). However, discrepancies between these databases have been acknowledged. Robeson et al. (8) found that Greengenes, SILVA, and GTDB presented sequence similarities but were taxonomically different, leading to a low proportion of taxonomic labelling shared among databases at all ranks below the domain level. Moreover, outlier sequences were found in the length distribution across databases, probably corresponding to partial or untrimmed 16S sequences, which are recommended to be discarded to avoid biases in the analysis. Additionally, SILVA and Greengenes exhibited an immense amount of unannotated or unknown labelled sequences at genus and species level (∼80%), which might introduce taxonomic noise during assignment (8).

To overcome these limitations and enhance classification performance, we created the GSR database for bacterial and archaeal 16S-based taxonomic profiling by integrating and manually curating the Greengenes, SILVA, and RDP databases. The taxonomic nomenclature of the GSR database has been unified to guarantee the coherence of annotations. Its performance has been compared with Greengenes, GTDB, SILVA, and RDP databases and other existing integrated databases, including ITGDB (9) and MetaSquare (10). The GSR database is available for full-length 16S sequences and the most commonly used hypervariable regions: V4, V1-V3, V3-V4, and V3-V5. It can be downloaded from the link (https://manichanh.vhir.org/gsrdb/).

## MATERIAL AND METHODS

### Creation of GSR database

#### Creation of the GSR full-16S database

A full-length 16S database, the GSR-DB (Greengenes-SILVA-RDP database), was created by merging three already existing databases: Greengenes (version 13_8, 99%) (4), SILVA (version 138, 99%) (6), and RDP (train set no.18) (7). A dataset with vaginal-related species was also included to ensure species detection for vaginal samples. The number of original entries of the Greengenes, SILVA, and RDP was 203,452, 436,681, and 21,194, respectively. Before the integration, taxonomy filtering and formatting were performed on each original database. Only Bacteria and Archaea kingdoms were retained from the databases, excluding Eukaryota and Virus kingdoms in the SILVA database. Additionally, a manual curation process was applied to ensure the removal of potential redundancies for the subsequent merging of the original databases. After this process, the percentage of retained entries was 10.05%, 17.08%, and 95.08% for Greengenes, SILVA, and RDP, respectively. The vaginal dataset was created with the 16S NCBI sequences proposed by Fettweis et al. (11) to create a vaginal reference database. Sequences and nomenclature corresponding to GenBank ids provided in the study were retrieved from the NCBI. Sequences without exact species names or those corresponding to non-16S sequences were excluded. Once all original databases were preprocessed correctly, they were merged using the forthcoming algorithm.

#### Manual curation of the GSR-DB

The curation process for the GSR-DB included several steps to ensure the quality and accuracy of the data. These steps involved manual identification and removal of patterns associated with unknown species (entries unannotated or with unknown labels, such as “uncultured”, “unidentified”, and “candidate”). Additionally, sequences that only provide information at the kingdom and species levels were discarded, particularly if they refer to rare bacteria from non-characterized environments (e.g. k_Bacteria,…, s_*bacterium_Te63R*.). Lastly, taxonomic nomenclature was carefully reviewed during the integration of databases, using the python module ETE toolkit (version 3.0) (12) to retrieve synonyms from the NCBI database. The NCBI taxonomy database (13) was chosen as the reference for taxonomic annotation as it enables the identification of synonyms for all the taxonomic annotations in the databases, and provides a standardised nomenclature. This procedure is capable of ensuring consistency and identifying misannotated organisms. One specific example mentioned is the identification of misannotated entries from the SILVA database, where certain entries labelled as bacteria are actually eukaryotic species, such as the annotation d_Bacteria; p_Proteobacteria; c_Gammaproteobacteria; o_Burkholderiales; f_Comamonadaceae; g_Paucibacter; s_*Cenchrus_americanus*, which is a plant species. This suggests that thorough steps were taken to ensure the accuracy and reliability of taxonomic information in the GSR-DB.

#### Merging algorithm

The algorithm used to merge the Greengenes, SILVA, RDP, and vaginal processed databases was based on the integration algorithms proposed by Hsieh et al. (9). The algorithm took two databases as inputs and integrated them as follows (**Figure 1**). First, one database was assigned as the reference database (R) and the other as the candidate database (C). Then, for each entry in the candidate database, it checked whether the candidate taxon (T_C_) was already present in the reference database. The candidate entry (sequence and taxon) was added to the dataset if not present. If T_C_ was present, the algorithm compared the candidate sequence (S_C_) to all the sequences in the reference dataset (S_R_) with the same nomenclature as T_C_. No integration was performed if S_C_ was identical or present as a substring in any of the S_R_. On the other hand, if S_C_ was not found in the reference dataset, the candidate entry (taxon and sequence) was added to the dataset. The RDP dataset was chosen as the first reference dataset for its taxonomic consistency, then the remaining datasets were added in the following order: SILVA, Greengenes, and vaginal (**Figure 2**). The resulting dataset is the GSR full-16S database.

**Figure 1.**
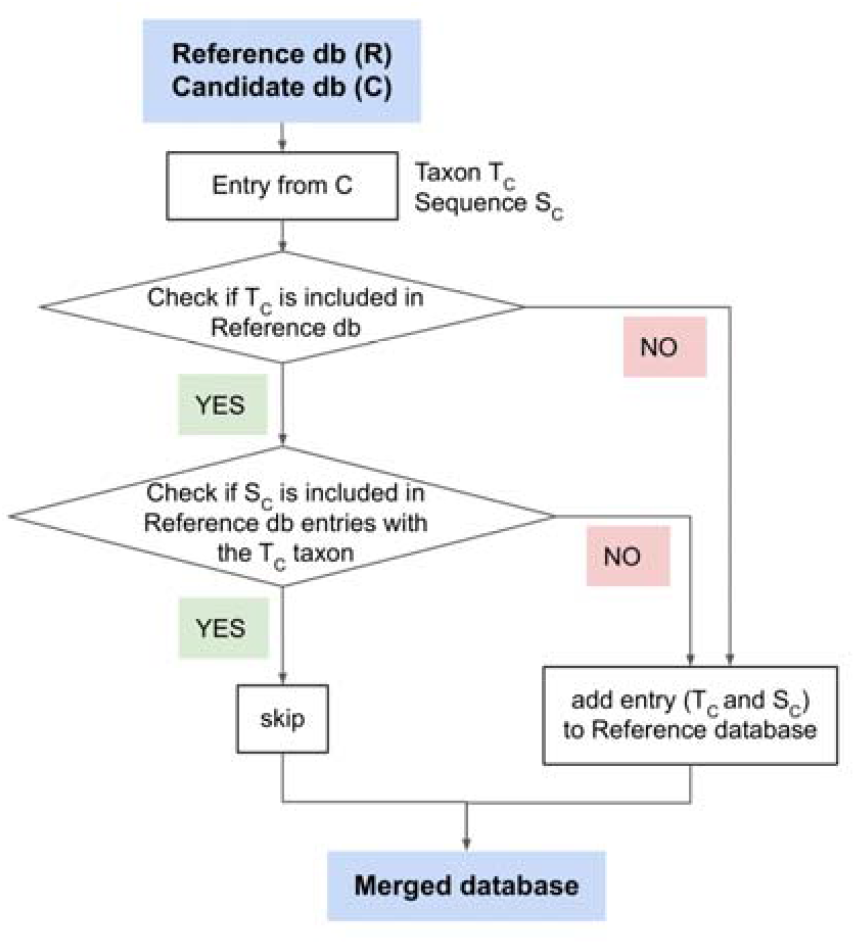
Merging algorithm to create the GSR database. The algorithm takes as an input a Reference database (R) and a Candidate database (C). Entries from the Candidate database are susceptible to be added to the Reference database after being evaluated.

**Figure 2.**
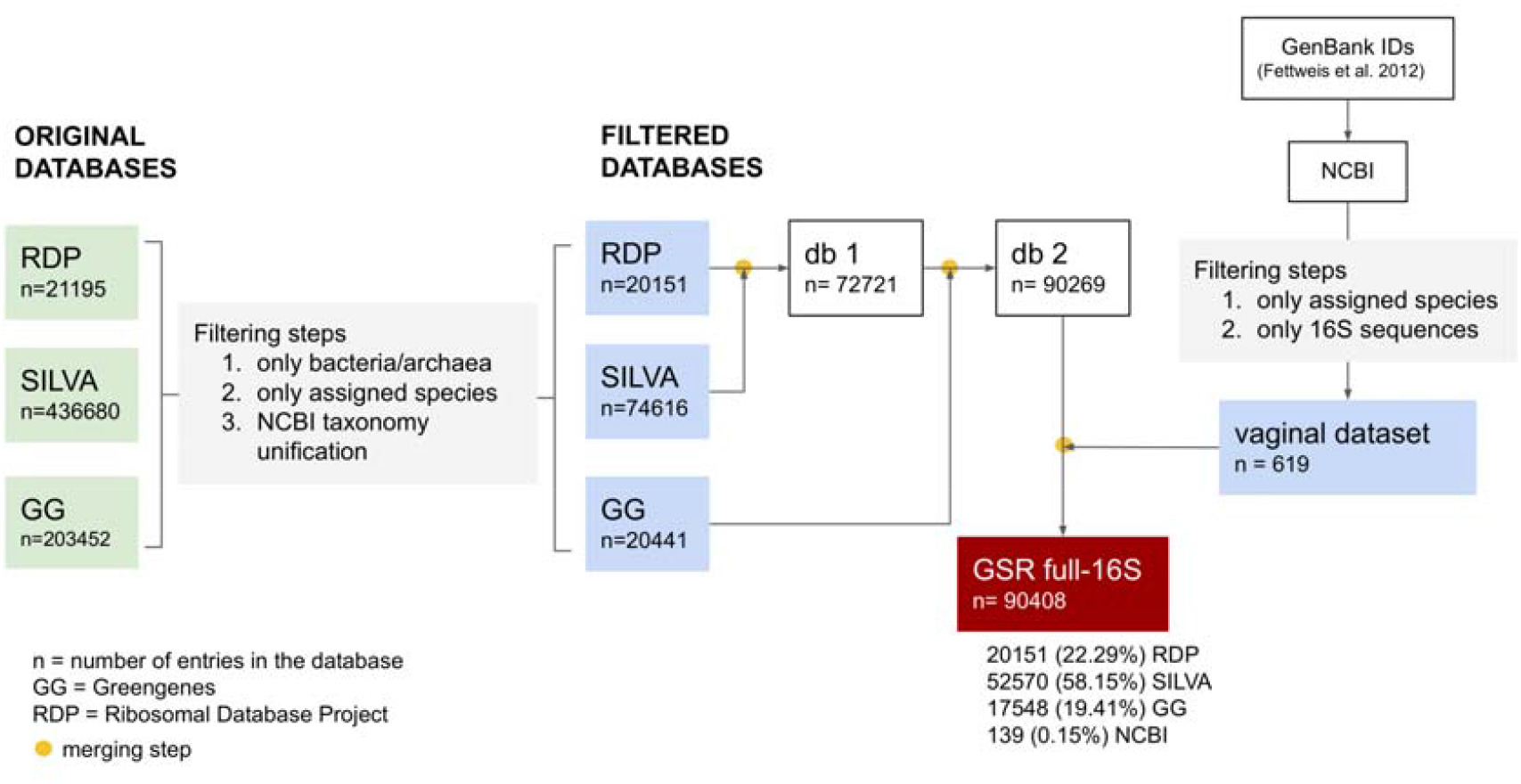
Database merging workflow to obtain the GSR full-length 16S database. It has a final size of 90408 entries with the following source composition: 22.29% RDP, 58.15% SILVA, 19.41% Greengenes, 0.15% NCBI. Merging steps were performed using the merging algorithm described in **Figure 1**.

### Creation of 16S variable region databases

#### Variable region extraction

The full-length GSR 16S database was used to create region-specific databases containing the most commonly used hypervariable regions in 16S analysis (V4, V3-V4, V1-V3, and V3-V5). The sequences for each region were extracted from the full-length GSR 16S database using the extract-reads function implemented in the QIIME 2 feature-classifier plugin (14) and the corresponding primers (3). QIIME 2 RESCRIPt plugin (8) was also used to dereplicate the resulting databases to remove redundant entries. These steps were also performed on the databases (Greengenes, GTDB, ITGDB, SILVA, and RDP) used in the validation analysis step.

#### Clustering

The region-specific GSR databases were clustered using CD-HIT (15,16) at 100% identity to host unique sequences and improve species detection. Nomenclatures of clustered sequences were merged into a single taxonomic name, as shown in the following example:

#### Nomenclatures to be clustered

1. *k__Bacteria; p__Firmicutes; c__Bacilli; o__Lactobacillales; f__Lactobacillaceae; g__Limosilactobacillus; s__Limosilactobacillus_fermentum*
2. *k__Bacteria; p__Firmicutes; c__Bacilli; o__Lactobacillales; f__Lactobacillaceae; g__Limosilactobacillus; s__Limosilactobacillus_oris*
3. *k__Bacteria; p__Firmicutes; c__Bacilli; o__Lactobacillales; f__Lactobacillaceae; g__Lactobacilus; s__Lactobacillus_crispatus*

#### Clustered nomenclature

*k__Bacteria; p__Firmicutes; c__Bacilli; o__Lactobacillales; f__Lactobacillaceae; g__Lactobacillus-Limosilactobacillus; s__Lactobacillus_crispatus:Limosilactobacillus_fermentum-oris*

### *In silico* mock community datasets

To assess the performance of the newly built database, three different mock communities (mockrobiota, vagimock, and gutmock) were constructed *in silico*. The vagimock and gutmock datasets simulate the relative abundance and species of biological samples from two different body sites. They were built from our GSR database with species commonly found in the human vagina and gut. The mockrobiota datasets were constructed using sequences obtained from the mockrobiota repository, a public resource for microbiome Bioinformatics benchmarking. From this repository, we recovered full-length 16S sequences provided only by datasets 3, 4, 5, and 12 to 23 of this repository (17).

Each in silico mock community dataset contained five samples, with given microbial abundance profiles, taxonomic and sequence information. The taxonomic information of the sequences was unified using the ncbi_taxonomy python module from the ETE toolkit (version 3.0) (12). Each mock community has a different level of complexity, which is crucial to reveal possible database issues (3). The composition of the mock communities at the species level can be found in **Supplementary Table 1**.

### Validation

#### Validation datasets

To evaluate the classification performance of the full-length and region-specific databases, we adapted the sequence length of the mock communities accordingly. For regions shorter than 460 nt (V3-V4 and V4), MiSeq Illumina reads were simulated using ART (18), using paired-end and single-end for V3-V4 and V4 regions, respectively. The corresponding parameters for each region were: V3-V4) -ss ‘MSv1’ –amp-na -nf 0 -l 250 -c 1 -rs 123 --minQ 25 -p -m 450; V4) -ss ‘MSv1’ -amp -na -nf 0 -l 200 -c 1 - rs 123 --minQ 25. Subsequently, the DADA2 module, (19) implemented in QIIME2, was used to denoise the reads and construct representative sequences (rep-seqs). To recover the expected nomenclature of rep-seqs, rep-seq sequences were mapped to their corresponding community sequences using a convolution method, as performed in the TAX CREdiT framework (14). For regions larger than 460 nt (full-16S, V1-V3, and V3-V5), sequences were directly treated as rep-seqs, due to software restrictions in simulating MiSeq (or PacBio) reads larger than 250 nt.

#### Classifier training and taxonomy assignment

It is known that some classifiers are strongly affected by parameter configurations. Therefore, different parameters for classifier training and taxonomy assignment steps were tested to find the optimal configuration. The sequences and taxonomy of each database were used to train the multinomial Naive Bayes (NB) Classifier implemented in q2-feature-classifier QIIME2 module (14). During this training, the n-gram-range parameter was tested with the values [6,6] and [7,7] (default), as its developers have already reported these ranges as optimal. Then, these classifiers were used to perform the taxonomy assignment of rep-seqs for each region. During the taxonomy assignment, the confidence threshold for limiting taxonomic depth was tested with the values “disable”, 0.5, 0.7 (default), 0.9, and 0.98. Evaluating two n-gram-range values and five confidence thresholds generated 10 different taxonomic profiling for each database.

#### Parameter comparison

To compare the performance of the 10 possible parameter configurations for each database, we calculated the average F1-scores across mock communities for each taxonomic level. The configuration with higher scores was retained for subsequent benchmarking of the databases.

#### Database benchmarking

The performance of the GSR database was compared with widely-used databases such as Greengenes, GTDB, SILVA, and RDP, but also with other available databases, such as ITGDB. Two independent approaches were used to assess the performances: the multi-class confusion matrix and the Bray-Curtis distances. The multi-class confusion matrix was used to evaluate the performance of a Machine Learning classification (e.g. Naive Bayes classifier) by comparing the expected sequence taxonomy versus the classified (**Table 1**). This confusion matrix allowed us to obtain validation metrics such as accuracy, precision, recall and F1-score by using the following equations:

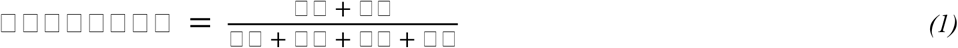

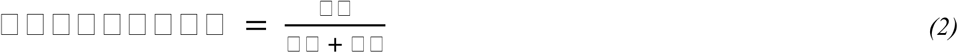

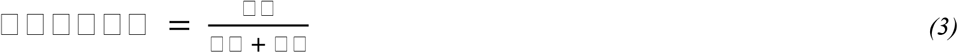

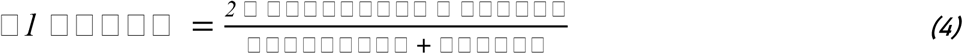

where TP is true positive, FP is false positive, TN is true negative, and FN is false negative.

The four metrics were measured at each taxonomic level as follows: A match was called when two taxonomic names (ID) were identical between the expected (E) and the assigned (A) name or, in the case of assignments with clustered databases, a match was called when one name was included in the other one (for instance, A = *Amylolactobacillus amylophilus* - *Lactobacillus iners*; E = *Lactobacillus iners*). For each taxonomic ID (T_i_): 1) TP was considered when T_i_ matched both A (assigned taxonomic ID) and E (expected taxonomic ID). 2) FP was defined when T_i_ matches A but not E. 3) FN was defined when T_i_ matches E but not A. 4) TN was defined when neither A nor E match T_i_.

**Table 1.** Example of a multi-class confusion matrix for Ti = *Lactobacillus iners*. TPs were all the *Lactobacillus iners* classified as *Lactobacillus iners*. FPs were all taxa classified as *Lactobacillus iners* that were not actual *Lactobacillus iners*. FNs were actual *Lactobacillus iners* not classified as *Lactobacillus iners*. TNs are other taxonomies different from *Lactobacillus iners* correctly classified as non-*Lactobacillus iners*.

|  |  | Expected Taxon (E) |  |  |
| --- | --- | --- | --- | --- |
|  |  | *Lactobacillus iners* | *Lactobacillus jensenii* | *Lactobacillus crispatus* |
| Assigned Taxon (A) | *Lactobacillus iners* | TP | FP |  |
|  | *Lactobacillus jensenii* | FN | TN |  |
|  | *Lactobacillus crispatus* |  |  |  |

Finally, validation metrics for all taxonomic ID were integrated using a weighted mean, taking the corresponding expected abundance as weight, using the following equation:

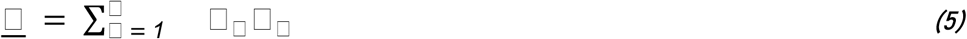

*<u>S</u>* = weighted mean score of the validation metric (precision, recall, f1-score or accuracy) for all taxonomies of a mock community

s_i_ = score of the validation metric for taxonomic ID T_i_

a_i_ = expected relative abundance of taxonomic ID T_i_ (weight)

n = total number of taxonomic IDs included in the mock community

Bray-Curtis distances were calculated between the expected and assigned composition for each sample in R (version 4.2.1) using the vegan package (version 2.6-4).

To discover significant differences in performance metrics, F1-scores and Bray Curtis distances were compared among the GSR, ITGDB, and SILVA databases using wilcoxon test (p-values adjusted by the Benjamini-Hochberg method).

Additionally, since different databases might use different taxonomic nomenclature, in order to consider synonyms of scientific names as correct matches, taxonomy unification (ETE toolkit v.3.0) was applied to each taxonomic classification before comparisons.

### Gut and vaginal microbial datasets

To further validate our database performance, we performed a case study using actual biological data from human gut samples (20) and human vaginal samples (21). These datasets contained V4 amplicon sequences. Taxonomic assignments were performed in both datasets using QIIME2 naive-bayes feature-classifier, with the following V4 databases as reference: Greengenes, GSR, GTDB, ITGDB, MetaSquare, SILVA, and RDP. The n-gram-range parameter was set to [7, 7] and the confidence threshold to “disable”, as these were the parameters found to perform best in the validation step.

#### Computational benchmarking

Furthermore, we also tested the computational cost of obtaining a taxonomic profile with the QIIME2 naive-bayes classifier with each of the V4 reference databases employed in this case study. We measured the time and memory consumption of the classifier training and the taxonomic assignment processes. Time was measured with the python built-in time module, and memory consumption was tracked using the memory_profiler module. These analyses were run on a computer with an Intel Xeon Gold 6238 processor with 44 CPUs and 187GB of RAM, and Ubuntu 18.04.4. Classifier training was run with default settings. Taxonomy assignment was performed setting the confidence threshold to “disable” and using 10 threads.

## RESULTS

### GSR Database

To optimise the prokaryotic taxonomic assignment, we created the GSR database by integrating and manually curating Greengenes, SILVA, and RDP datasets (**Figures 1 and 2**). The integrated full-length 16S GSR database has a total size of 90408 sequences, with the following source composition: 22.29% RDP, 58.15% SILVA, 19.41% Greengenes, and 0.15% NCBI (vaginal-specific sequences). The source composition and total size of the variable region databases are shown in **Table 2**. The V1-V3 and V4 databases are those with fewer available sequences. The sequence length distributions of the GSR databases are presented in **Figure 3**.

**Table 2.** Source composition of GSR databases. Number of entries of each GSR database that were recovered from each source database.

| Database | Region | Cluster | Source databases |  |  |  | Total |
| --- | --- | --- | --- | --- | --- | --- | --- |
|  |  |  | RDP | SILVA | Greengenes | NCBI |  |
| GSR | V1-V3 | 100% | 6707 (24.96%) | 14595 (54.31%) | 5521 (20.54%) | 51 (0.19%) | 26874 |
|  | V3-V4 | 100% | 15401 (30.79%) | 22621 (45.23%) | 11949 (23.89%) | 45 (0.09%) | 50016 |
|  | V3-V5 | 100% | 16659 (29.43%) | 26239 (46.35%) | 13655 (24.12%) | 53 (0.09%) | 56606 |
|  | V4 | 100% | 12670 (32.65%) | 16186 (41.72%) | 9916 (25.56%) | 29 (0.07%) | 38801 |
|  | full-16S | none | 20151 (22.29%) | 52570 (58.15%) | 17548 (19.41%) | 139 (0.15%) | 90408 |

**Figure 3.**
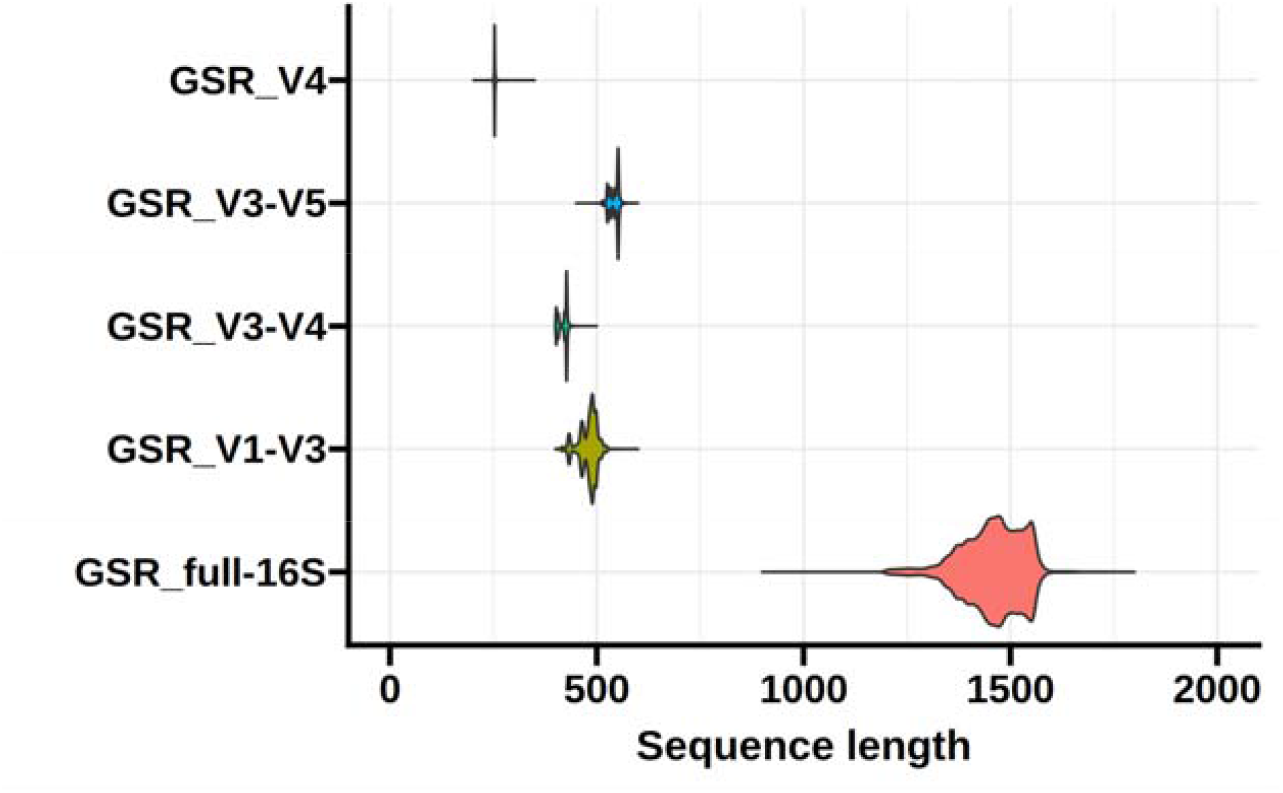
Sequence length distribution. Sequence length distribution of the GSR databases. Databases for the variable regions are clustered at 100% identity.

### QIIME2 parameters impact taxonomic assignment performance

The 16S rRNA analysis pipeline of QIIME2 (22) includes training a Naive-Bayes classifier with a reference database and a subsequent taxonomic assignment of the rep-seqs (14). In these two steps, we tested different values of n-gram-range and confidence threshold parameters for each database (full-length 16S and specific 16S regions) as it is known to affect the classifier’s performance. **Figure 4** summarises the performance of the aforementioned parameters across tested databases and regions. Two n-gram-range values were tested: [6,6] and [7,7]. The Wilcoxon test showed that [7,7] performed better than [6,6] (p<0.0001) (**Figure 4A**). Confidence threshold values show significant differences in F1-score (**Figure 4B**, p<0.0001 for all comparisons in a pairwise manner), precision, and recall at both genus and species levels (**Supplementary Tables 2, 3, and 4**). Setting the confidence threshold to ‘disable’ provided the best classification results at the species level, suggesting that setting a confidence threshold for the QIIME2 classifier notably restricts the predictions at the species level without improving the predictions at higher levels. Therefore, the n-gram-range of [7,7] and ‘disable’ confidence threshold were further used to benchmark the GSR database with other already existing databases.

**Figure 4.**
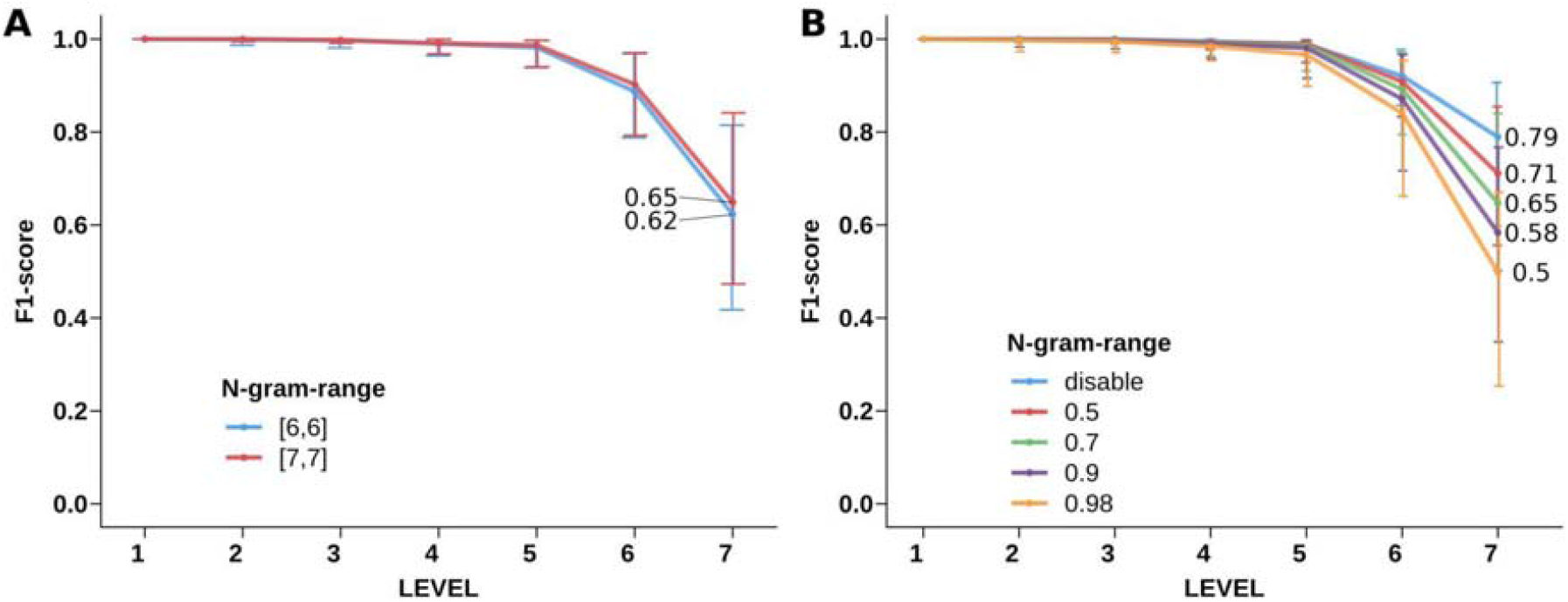
Benchmarking of n-gram-range (A) and confidence threshold (B). The median F1-score is shown at each taxonomic level across all tested databases, regions, and validation datasets. Error bars represent the interquartile range (N=900). Full data for the family, genus, and species level is available in Supplementary Tables 2, 3, and 4.

### GSR outperforms existing databases across all tested regions

To assess the performance of the newly created database, we benchmarked the GSR database with the other existing databases (Greengenes, SILVA, RDP, ITGDB, MetaSquare), using two different approaches: validation metrics (F1-score shown in **Figure 5**, precision and recall shown in **Supplementary Tables 5, 6**, and **7**) and Bray-Curtis distances (**Figure 6, Supplementary Table 8**). In order to increase the robustness of the results, we defined the combination of the F1-score and the Bray-Curtis distance as the validation scores. The database with the best validation scores will achieve the highest F1-score and the shortest Bray-Curtis distance.

**Figure 5.**
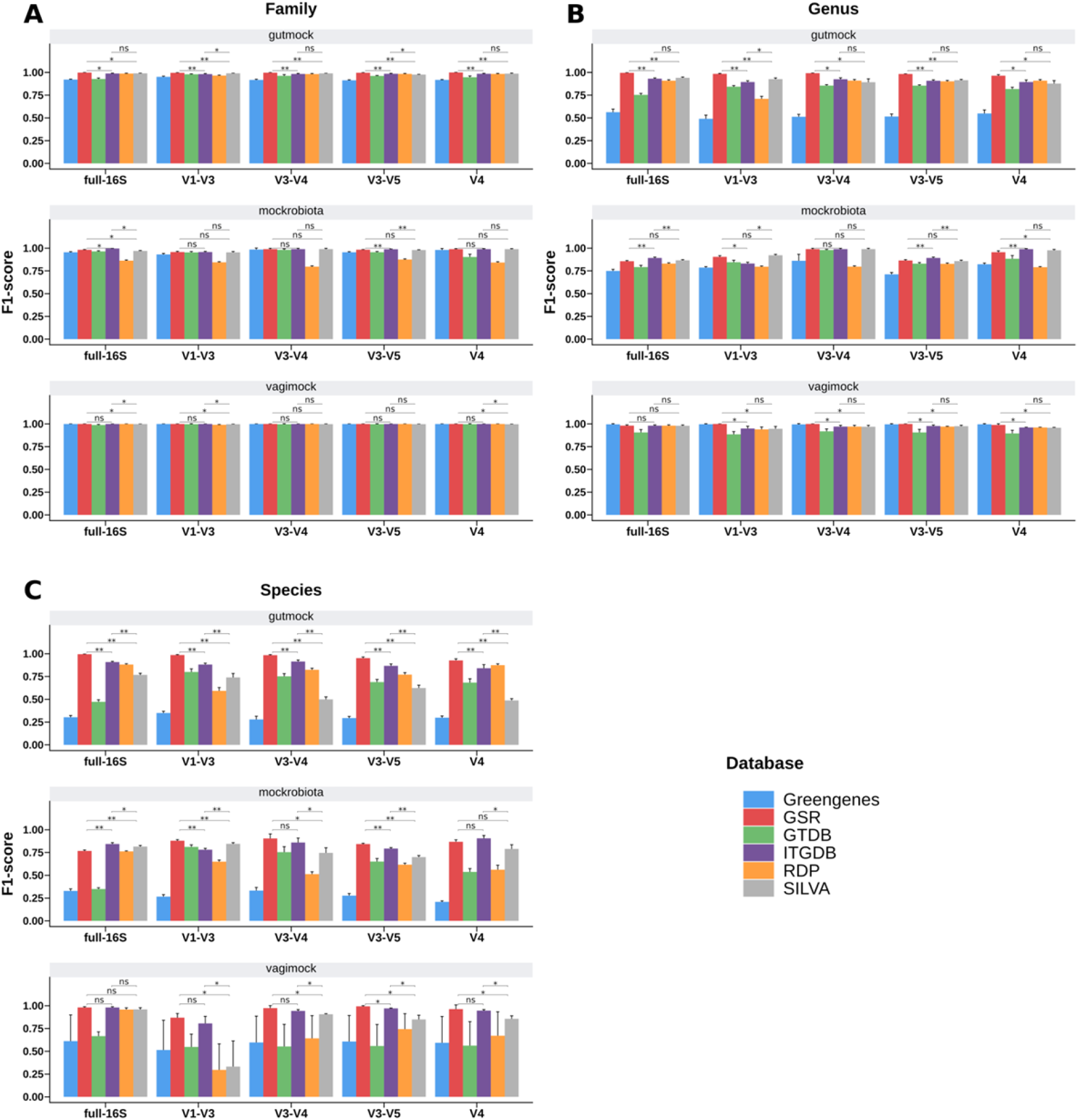
Database benchmarking at family (A), genus (B), and species (C) levels using validation metrics. The mean F1-score across the five metagenomic samples is shown for each evaluated region and dataset. Error bars are the standard deviation. Precision and recall metrics are available in Supplementary Tables 5, 6, and 7 for family, genus, and species levels, respectively. Wilcoxon test was conducted between F1-scores of GSR, ITGDB, and SILVA databases. ns=not significant; * = padj < 0.05; **= padj < 0.01; ***= padj < 0.001.

**Figure 6.**
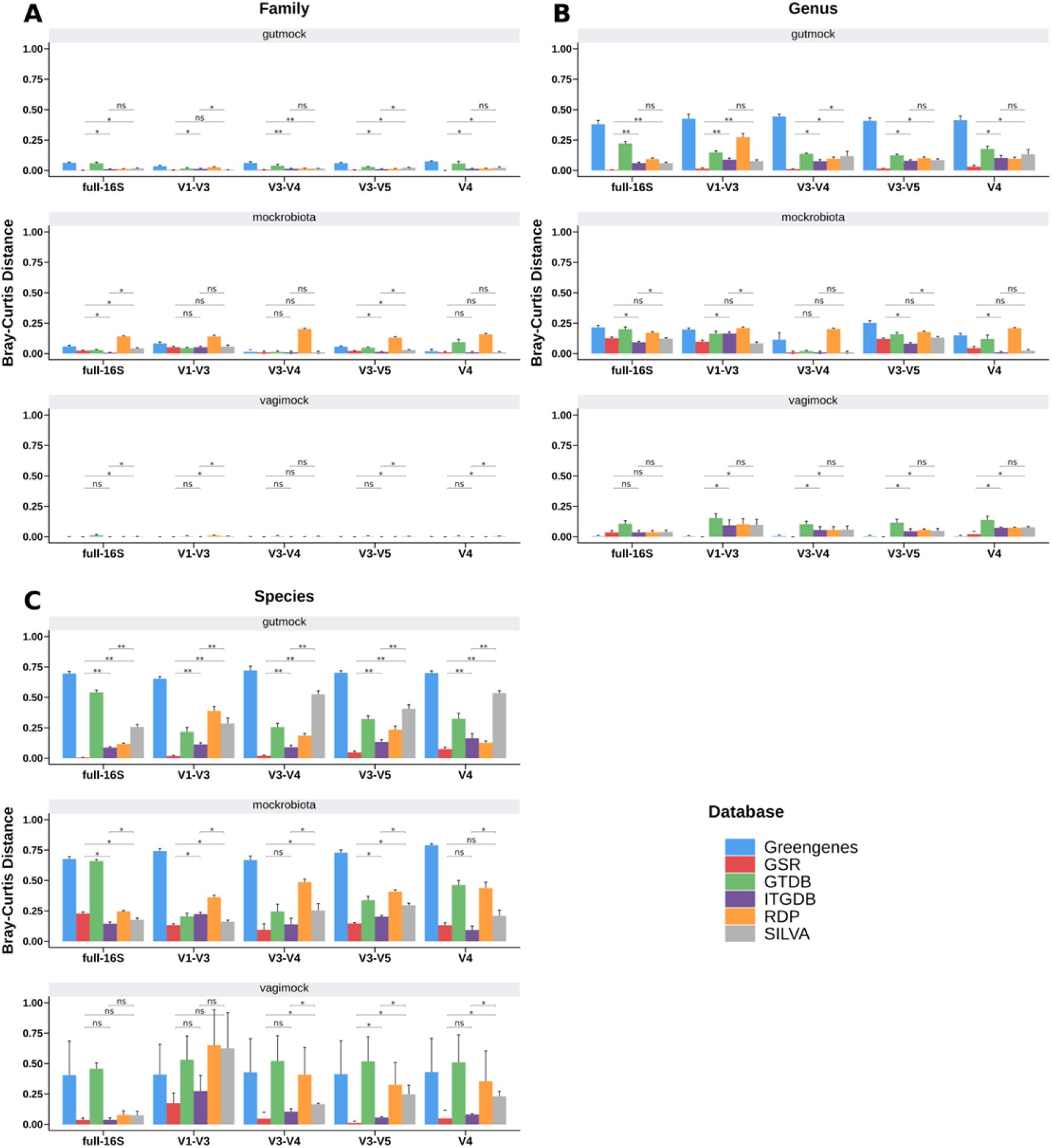
Database benchmarking using Bray-Curtis distances between expected and observed composition at family (A), genus (B), and species (C) levels. The mean Bray-Curtis distance across the five metagenomic samples is shown for each evaluated region and dataset. Error bars are the standard deviations. Data is available in Supplementary Table 8. Wilcoxon test was conducted between F1-scores of GSR, ITGDB, and SILVA databases. ns=not significant; * = padj < 0.05 ; **= padj < 0.01; ***= padj < 0.001.

At family level, the GSR, ITGDB and SILVA databases reached in most cases the best validation scores across regions in all the validation datasets (**Figure 5A, Figure 6A, Supplementary Tables 5 and 8**). At genus level (Fig5B, Fig6B, Supplementary table 6 and 8), GSR achieved significantly better validation scores across almost all regions in the gutmock dataset, followed by ITGDB and SILVA databases. In the mockrobiota dataset, ITGDB and SILVA achieved the best validation scores across regions, sharing similar values with GSR and for V1-V3 and V3-V4. Finally and most importantly, at the species level (**Figure 5C, Figure 6C, Supplementary Tables 7** and **8**), except for the full-16S region where ITGDB had the best validation score, the GSR database presented the best scores for almost all the regions and validation datasets. Overall, these results indicate that whereas the database performance remains relatively stable up to the family level, substantial differences were observed at the genus and species performance, with GSR showing the best performance results among the tested databases.

The Greengenes database performed worst in all tested environments, except for the vagimock dataset at the genus level, for which it performed similarly to the other databases. On the other hand, the RDP, GTDB, and SILVA databases yielded better results across all environments and regions. Previous studies have already pointed out the increased accuracy of SILVA and RDP databases in comparison to Greengenes (3), mainly due to the fact that, in the last few years, SILVA and RDP have been updated more frequently than Greengenes. The better performance of Greengenes in identifying genus-level classifications within the vagimock dataset could be attributed to the low complexity of this mock community. It has been observed that database limitations may not be as apparent when analysing mock communities with limited characteristics (3).

### Case study: vaginal and gut datasets

In methodological benchmarking studies, it is crucial to contextualise the benchmarking outcomes using actual biological data. Therefore, 10 vaginal and 10 gut microbiome samples were analysed from Vargas et al., 2022 (21) and Yáñez et al., 2021 datasets, containing 2089 and 31885 V4 representative sequences, respectively. These datasets allowed us to assess the consistency of taxonomic nomenclature among our newly built database and other existing databases, including Greengenes, GTDB, ITGDB, RDP, and SILVA. Additionally, the analysis of real datasets allowed us to compare the computational cost of taxonomy profiling among the aforementioned reference databases.

#### GSR annotation enhances taxonomic nomenclature consistency

Each database uses different synonym terms for the same NCBI taxonomy ID, as shown in **Figure 7**. For instance, in **Figure 7A**, Greengenes, GSR, and RDP use exclusively the term *Bacteroidetes* for NCBI:txid976, SILVA and GTDB use the synonym *Bacteroidota* and ITGDB uses both of them. Similarly, for NCBI:txid201174, Greengenes, GSR and RDP use the term *Actinobacteria*, SILVA and GTDB use the synonym *Actinobacteriota* and ITGDB uses both of them. In addition, GTDB splits the phylum *Firmicutes* into several clusters, namely *Firmicutes, Firmicutes_A, Firmicutes_B, and Firmicutes_C*. At the order level, another example can be found in **Figure 7B**. For NCBI:txid186802, Greengenes and RDP use the term *Clostridiales*, and GSR uses the synonym *Eubacteriales*. SILVA, GTDB, and ITGDB use several non-NCBI terms such as *Clostridia, Lachnospirales*, and Oscillopirales. Moreover, ITGDB also uses the accepted term *Clostridiales*. Finally, other taxonomy inconsistencies can be found at the family level (**Figure 7C**). For NCBI:txid216572, whereas Greengenes uses the term *Ruminococcaceae* and GSR uses its synonym *Oscillospiraceae*, SILVA, GTDB and ITGDB use both aforementioned terms, and SILVA and ITGDB also use the synonym *Hungateiclostridiaceae*. Taken together, these results indicate that Greengenes, RDP and GSR databases have robust taxonomic nomenclatures, using exclusive terms for one NCBI taxonomy id. In contrast, SILVA, GTDB, and ITGDB databases use several terms to refer to the same taxon, some of which are non-NCBI terms.

**Figure 7.**
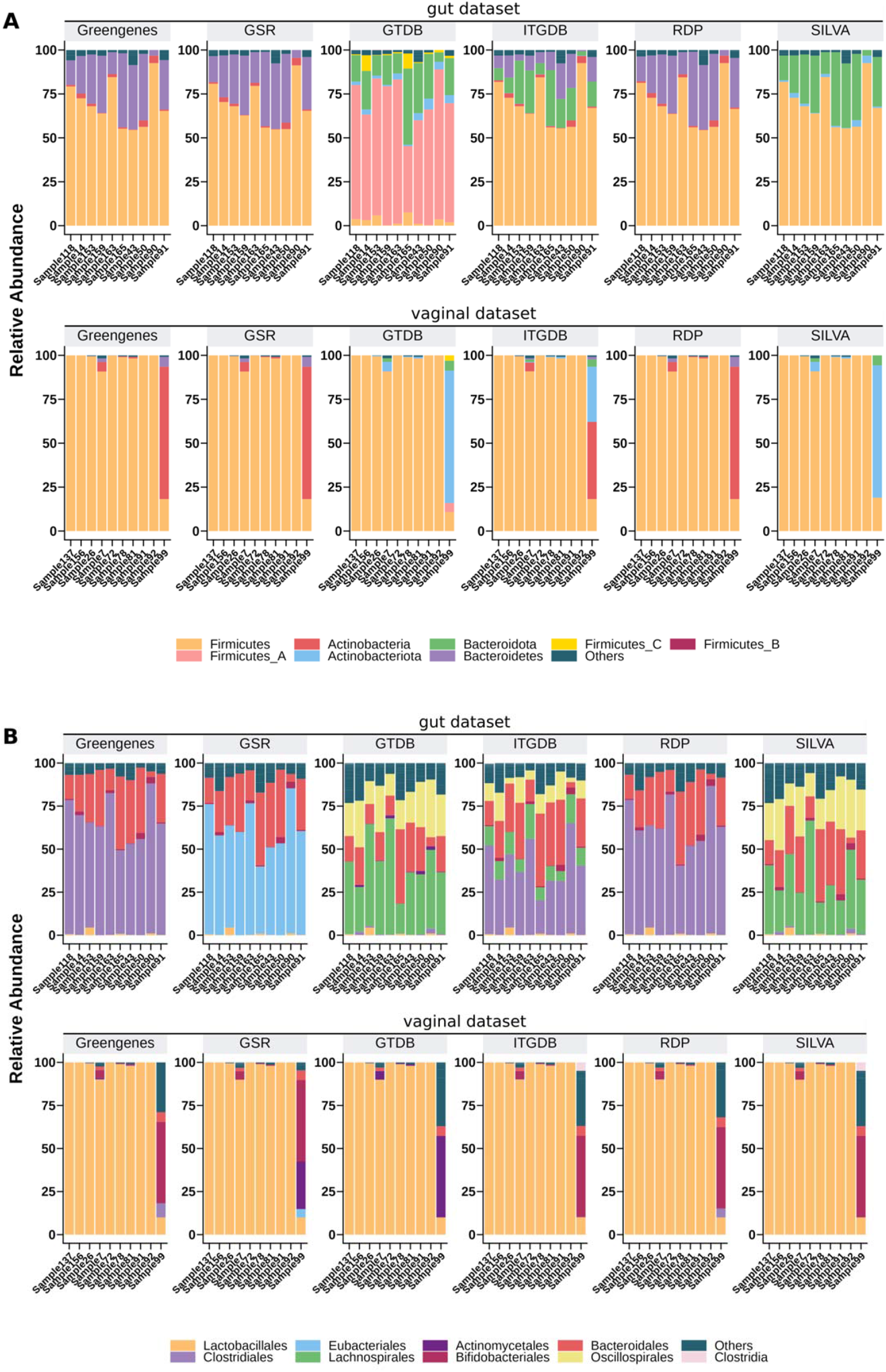
Relative abundance of gut and vaginal samples at phylum (A) and order (B) levels. Only relevant taxa are displayed. The remaining taxa are included in the label ‘Others.’ Family level is available in Supplementary Figure 1.

#### Computational benchmarking

The benchmarking was performed in two steps in which the databases were involved: classifier training and taxonomic assignment. A Naive Bayes classifier was trained in QIIME2 using the V4 region of each reference database. Training time and memory usage for each classifier are shown in **Table 3**. The most computationally efficient classifier training was obtained using the RDP, GSR, or Greengenes databases. These three classifiers were trained within three minutes and required less than 7GB of RAM. ITGDB and GTDB classifiers show an increased computational cost, doubling the time and memory usage of the aforementioned ones. The SILVA classifier required a significantly higher amount of computational resources, taking up to 40 minutes and 25 GB of RAM to be trained. The MetaSquare classifier was the most computationally expensive to train, being time-consuming and memory intensive.

**Table 3.**
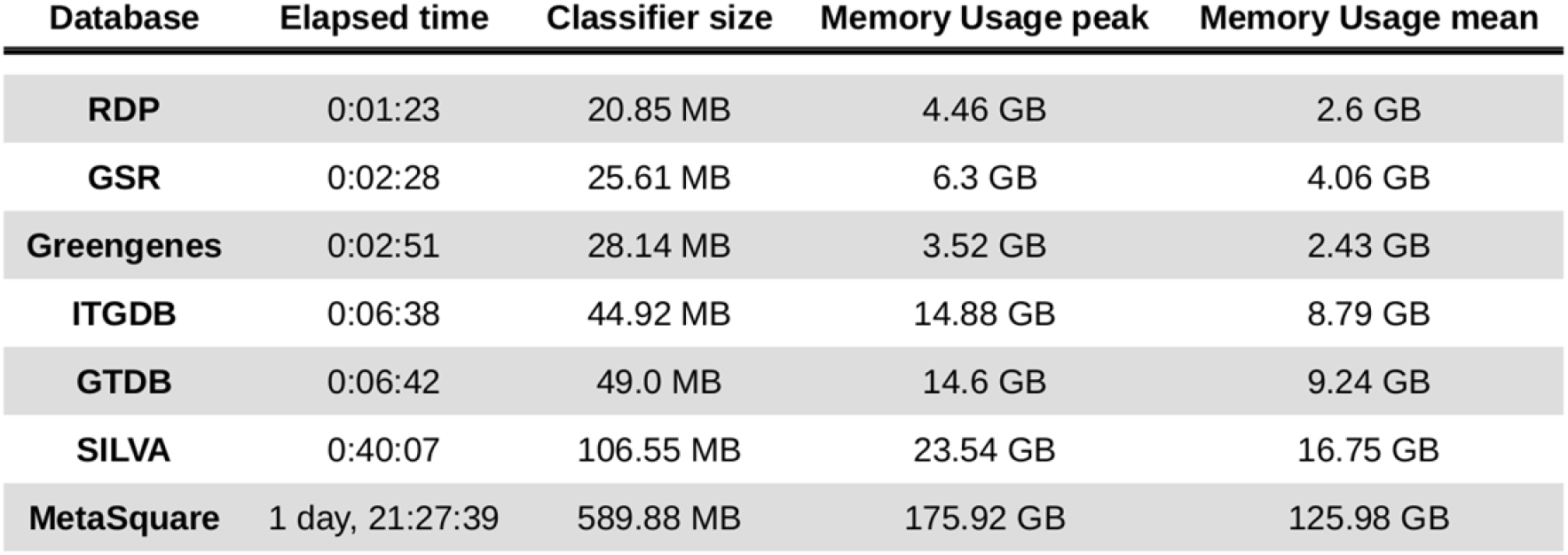
Benchmarking results for the classifier training step. A QIIME2 naive-bayes classifier was trained with each one of the reference databases using the default parameters.

The trained classifiers were then used to perform a taxonomy profiling of the intestinal and vaginal datasets, with the confidence threshold set to ‘disable’ and multithreading used with 10 threads. The benchmarking results for this step are presented in **Table 4**. Resource consumption resembled the pattern seen in the classifier training step. Greengenes classifier was the most computationally inexpensive, followed by GSR and RDP, which are still affordable. ITGDB and GTDB almost double the required resources, and SILVA was the most resource-consuming. MetaSquare classifier was also tested, but its taxonomy assignment could not be completed due to a lack of computational resources.

**Table 4.** Benchmarking results for the taxonomy assignment step. Gut and vaginal datasets were taxonomically profiled using previously trained classifiers of each reference database. MetaSquare classifier was also tested but no results were obtained due to a lack of computational resources (> 187 GB RAM).

| Dataset | Database | Elapsed time | Memory Usage peak | Memory Usage mean |
| --- | --- | --- | --- | --- |
| gut | Greengenes | 0:00:17 | 7.09 GB | 3.11 GB |
|  | RDP | 0:00:46 | 18.21 GB | 6.97 GB |
|  | GSR | 0:01:08 | 18.55 GB | 5.11 GB |
|  | ITGDB | 0:01:28 | 33.62 GB | 11.74 GB |
|  | GTDB | 0:01:44 | 37.47 GB | 13.46 GB |
|  | SILVA | 0:02:17 | 48.93 GB | 16.36 GB |
| vagina | Greengenes | 0:00:09 | 5.64 GB | 2.55 GB |
|  | GSR | 0:00:29 | 14.85 GB | 4.33 GB |
|  | RDP | 0:00:29 | 12.56 GB | 4.28 GB |
|  | ITGDB | 0:01:00 | 26.59 GB | 6.83 GB |
|  | GTDB | 0:01:07 | 30.12 GB | 7.83 GB |
|  | SILVA | 0:01:30 | 37.86 GB | 9.28 GB |

## DISCUSSION

In this study, we generated a new 16S database for prokaryotic and archaea organisms: the GSR database. The performance of the GSR database was assessed in conjunction with five other existing 16S databases: Greengenes, GTDB, ITGDB, SILVA, and RDP. Our attempt to evaluate the MetaSquare database was hampered due to its extremely high demand for computational resources compared with other existing databases (**Table 3**). We believe these requirements are unreasonable and impractical. Therefore, we discarded MetaSquare for subsequent analysis and cannot rationally advise its use.

Before database comparisons, we first explored the best parameter configuration for each of the five databases, as previous studies have pointed out the impact of n-gram-range and confidence threshold parameters on classifier performances (14). The n-gram–range value of [7,7] performed better than [6,6], whereas the confidence threshold value of “disable” significantly outperformed at the species level. These results suggest that confidence threshold value plays an essential role in the taxonomic resolution and should be consistently reported in studies. Based on our results, we recommend using the n-gram-range of [7,7] and confidence threshold “disable” values in microbial profiling studies that utilise the GSR database.

Regarding database performance, GSR outperformed GTDB, SILVA, RDP, and Greengenes databases in almost all tested environments and regions. ITGDB database presented a comparable performance to the GSR database, performing better in the mockrobiota dataset. However, the ITGDB database has some significant shortcomings, not detected in the GSR database, such as taxonomical discrepancies and lower computational efficiency.

The case study performed with gut and vaginal sample datasets exposed the consequences of not unifying the taxonomy when merging databases with different taxonomic annotations. ITGDB database presented multiple cases of taxonomical inconsistencies (**Figure 7**), where several synonym terms were used to refer to the same taxonomic clade. A similar behaviour is also noticeable in SILVA but to a lesser extent. The lack of a consistent taxonomy might severely interfere with microbial taxonomy analyses, impacting diversity metrics or differential composition analyses. In this regard, it is worth noting that the GSR database does not suffer from taxonomic consistency issues and can provide more reliable and robust results. Furthermore, this case study revealed that the computational resources used by QIIME2 differ depending on which reference database is employed. ITGDB database made QIIME2 consume twice as many computational resources as GSR (**Tables 3 and 4**), making GSR a more suitable alternative for obtaining high-resolution taxonomy profiles at lower computing costs.

Despite the described results, several limitations need to be considered. Firstly, the lack of testing on non-human samples, such as soil and water samples, raises concerns about the generalisability of the database to different contexts. Without this information, we cannot fully understand how well our database will perform in these environments. Secondly, the GSR database only contains sequences from known species, excluding unclassified organisms or organisms labelled as uncultured. While this may be detrimental for the analysis of environments containing a large amount of unknown or uncultured species (8), we demonstrated that it improves species detection in well-described environments, such as human body sites. Additionally, the utilisation of a single classification software (QIIME2 naive bayes) precludes the ability to extrapolate the performance of our database to alternative classification methods, as the use of different software may yield different results. Finally, another limitation is the restricted testing conducted in human-like environments. Although gut and vagina samples have been examined in this study, the database usefulness could be more comprehensively evaluated by extending the analysis to other human environments, such as skin and saliva. Overall, the GSR database demonstrates potential, but it is crucial to acknowledge and address the aforementioned factors to obtain a thorough understanding of its applications and potential drawbacks.

While 16S amplicon-based sequencing has limitations, its low cost and simplified methodology still make it a valuable tool for analysing the microbiome composition. The vast amount of data generated during the last decade cannot only help to answer pressing questions about microbiome-disease relationships in larger epidemiological studies, but also can be used along with shotgun metagenomic sequencing data to explore new clinical applications (23). Therefore, the GSR database offers several advantages for microbial taxonomic classification using 16S sequencing. It integrates three of the main reference databases, ensuring a comprehensive and accurate taxonomic annotation. The taxonomy consistency allows for reliable analysis, which is crucial for the robustness of microbiome studies. GSR database also demonstrates an improved performance with microbial communities containing mainly known species, enhancing its utility in various applications. Finally, its usage is not computationally expensive, making it accessible to researchers with limited computational resources. Overall, these features make the GSR database a valuable resource for the scientific community to further investigate microbial communities.

## DATA AVAILABILITY

The database and the validation data will be made available in the following link: https://manichanh.vhir.org/gsrdb/

## SUPPLEMENTARY DATA

Supplementary Data are available at NAR online.

## AUTHOR CONTRIBUTIONS

Leidy Alejandra G. Molano: Conceptualization, Formal analysis, Methodology, Software, Validation, Writing—original draft. Sara Vega-Abellaneda: Conceptualization, Formal analysis, Methodology, Software, Validation, Writing—original draft. Chaysavanh Manichanh: Conceptualization, Supervision, Funding acquisition, Writing—review & editing.

